# What Relative Neocortex Size Tells Us About Social Evolution

**DOI:** 10.1101/2025.11.26.690799

**Authors:** R.I.M. Dunbar

## Abstract

Primate evolution has been dominated above all by increases in brain size over time. Here I show that two separate pressures have been responsible for substantial changes in the major structural features of the primate brain. Lineages that have adopted pairliving are associated with a differential increase in the volume of the subcortical brain. This mainly reflects an increase in body size and a consequent need to invest in cerebellum size in order to manage large body masses in three-dimensional arboreal environments, a trend that continues through the great apes. The second trajectory is associated with a switch to group-living, and is associated with a progressive increase in neocortex volume. This seems to have occurred in three distinct waves corresponding to stepwise increases in social group size. These trends are not associated with phylogeny, but represent taxonomically mosaic evolution driven by individual species’ exposure to new kinds of habitats.

## Introduction

Mammalian evolutionary history has been dominated by changes in brain size, though some lineages (anthropoid primates, delphinids, tylopods, equids, canids) exhibit much stronger signals than others (felids, ruminant ungulates, ceratomorphs) where brain size has changed very little over geological time) (Shultz & Dunbar 2010). Within primates, these changes in brain size have largely involved the neocortex (which accounts for 50-80% of total brain size) (Finlay & Darlington 1995) and the evolution of stable, bonded social groups (Shultz & Dunbar 2010, 2022; Dunbar & Shultz 2021a, 2023). Neuroimaging studies in both humans and primates have focussed this relationship down onto the prefrontal cortex and the default mode neural network (DMN: the single largest connectome in the brain) (Andrews et al. 2010; Powell et al. 2012; Mars et al. 2012; Spreng et al. 2020). The DMN links processing units in the prefrontal cortex, the temporo-parietal junction and the temporal lobe, with important extensions down into the limbic system and the cerebellum.

More detailed analysis of primates has revealed that the social brain relationship is not formed by a single homogenous dataset; rather, it consists of a series of distinct socio-cognitive grades (Dunbar 1993; Dunbar & Shultz 2021a). These grades have little or no taxonomic signal, but instead represent the demography-dependent mosaic evolution of distinctive social styles related to the bonding processes needed to hold groups of different size together (Dunbar 2025). Species that live in smaller groups use simple attention mechanisms to hold the group together; those in medium-sized groups invest more heavily in social grooming to build a wider network of bonded relationships; while those that live in large groups have evolved specialised social cognitive skills that allow them to manage relationships with individuals they do not interact with regularly.

Here, I explore in more detail the relationship between social group size and relative investment in cortical versus subcortical neural tissue in primates. Since the DMN and its associated units occupy almost the entire neocortex other than relatively small regions associated with specialised functions (the association and motor cortices; the visual system in the occipital lobes), treating the neocortex as a functional unit is a reasonable approximation. This is fortunate, because no brain datasets subdivide the neocortex (aside from differentiating the visual system in the occipital lobe). My central question is whether there are grade differences in relative neocortex volume that correlate with species’ social styles. In primates, social style varies between strongly bonded groups (pairbonded monogamy, larger stable multimale/multifemale groups), smaller more casually bonded groups (typically harems) and large multilevel groupings based on fission-fusion dynamics. Having established that there grade differences, I ask whether these are associated with tradeoffs between neocortex volume and the volume of subcortical brain regions.

## Methods

Except for the orang utan, all brain volume data are from Stephan et al. (1981) since this is still the largest and taxonomically most diverse brain database that distinguishes between neocortex and subcortical regions. Brain data for the orang utan are from Bush & Almann (2004). Mean social group size for individual species are taken from the comprehensive dataset provided by Dunbar et al. (2018), except for *Miopithecus* for which I use the sleeping band (N=37.4) following Dunbar & Shultz (2021a) and *Pongo* for which, following Dunbar & Shultz (2021a, 2023), I use the resident community as defined by Singleton & van Schaik (2001) (see Dunbar & Shultz 2025). Although the mean community size for *Pan* is 42.5 (Dunbar et al. 2018), in fact the genus has a bimodal distribution with mean community sizes of 32 and 59 (independent of species identity). I therefore use both values for this genus. Diet data are from Powell et al. (2014) subject to the correction for *Macaca mulatta* introduced by Dunbar & Shultz (2025).

### The data are given in the *ESM*

I first regress neocortex volume on total brain volume. The two are highly correlated (OLS regression: Neocortex volume = -2.516 + 0.735*BrainVolume, with both volumes in cc; r^2^=0.998, β=0.999, F_1,43_=25628.1, p<<0.0001). Since r^2^>0.95, I calculate residuals of actual neocortex volume from this line to give an index of a species’ relative investment in cortical versus subcortical brain tissue. I then regress group size onto these residuals. Since, in this case, r^2^<<0.95, OLS underestimates the regression slope, with a degree of underestimate directly proportional to the poorness of the fit. This is because, in OLS regression (which was designed for use in experimental designs where the X-value is determined by the experimenter), the error variance (in the form of the residual on the Y-axis) forms part of the calculation of the slope. When there is error variance on both axes (and the data are hence bivariate uniform rather than bivariate normal as required by OLS), RMA regression (which places the regression line up the centre of the distribution) is recommended (Kendall & Stuart 1979; Rayner 1985). Because RMA regression minimises the error variance on both the X and Y axes, I determine residuals orthogonal (i.e. at right angles) to the regression line (Fig. 1).

**Fig. 1.**
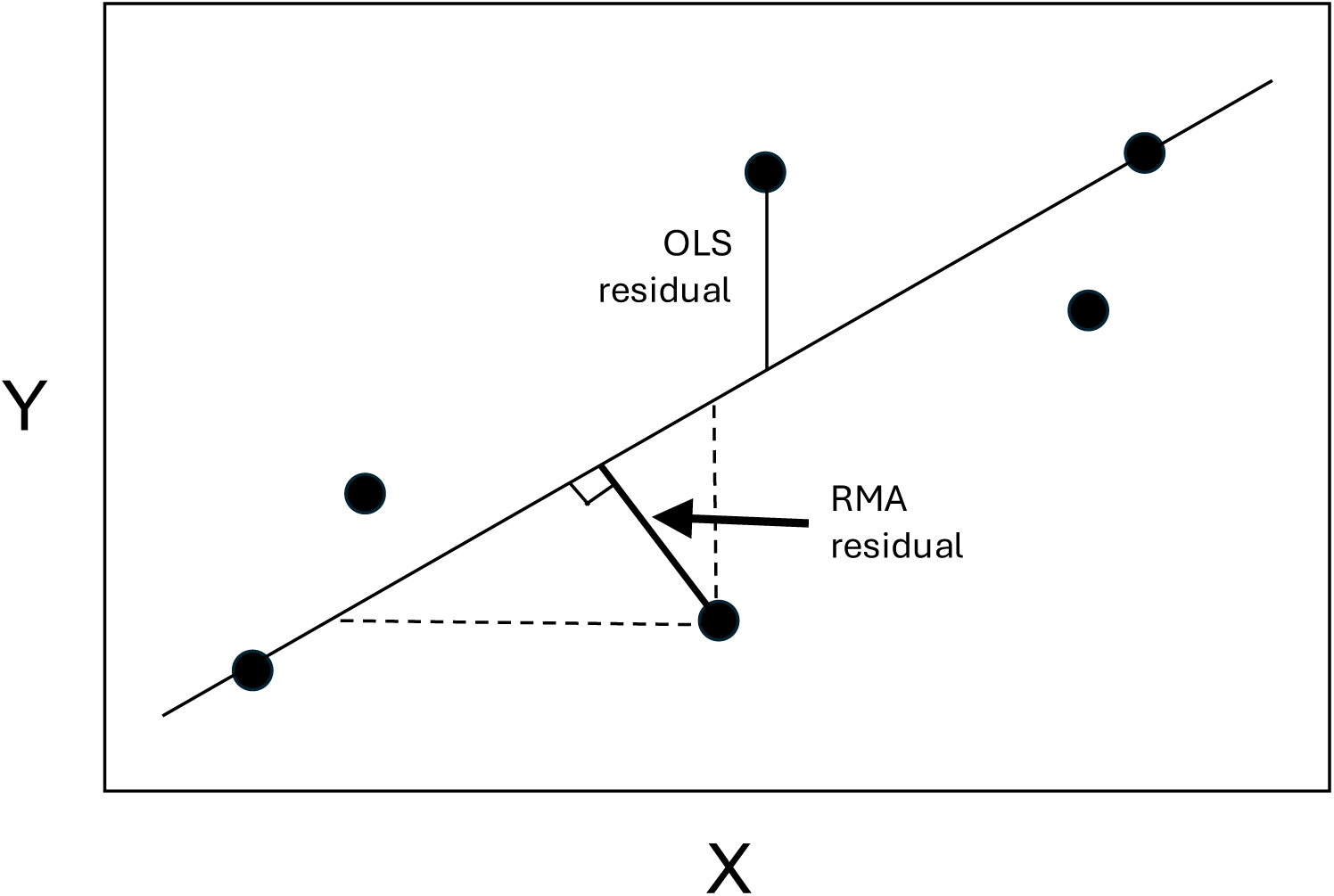
Calculating residuals for OLS and RMA regressions.

I use *k*-means cluster analysis on these residuals to determine the optimal number of distinct clusters within the range 2≤*k*≤10. The optimal number of clusters is that which maximises goodness-of-fit (here indexed by the F-ratio) while minimising the number of clusters, subject to the proviso that there should be as few clusters as possible with N=1. This can be identified by the inflexion point where the increase in goodness-of-fit as a function of number of clusters begins to asymptote. This is given by the point on the X-axis where the regression line is 1/e^th^ down from the asymptotic value on the Y-axis.

Kronmal (1993) cautions against the use of ratios in regression analyses on the grounds that we cannot tell whether the effect is due to a change in the numerator or denominator, or both (see also Deacon 1990). He recommended that the regression analysis be run with both components of the ratio as separate predictor variables (in the form Y= X + Z^-1^). To disentangle the effects of changes in numerator and denominator, I ran the regression analyses for both Z and Z^-1^ as well as X.

I do not use phylogenetic methods in these analyses for three reasons. First, in primates, the phylogenetic signal in socioecological and lifehistory indices is negligible (Kamilar & Cooper 2013), and especially so at genus level (Dunbar & Shultz 2021a, 2023). In some 30 tests of the relationship between group size and brain size in primates where the analysis was run with and without phylogenetic control, none has yielded a difference in outcome. As it happens, with just three exceptions, Stephen et al. (1981) sampled only one species in each genus. Autocorrelation between genera is inevitably much lower than that between species of the same genus. Second, we seek to determine whether grades exist in the data. All phylogenetic methods deliberately smooth over these effects in order to test for an overall relationship. More importantly, they have difficulty dealing with grades unless these are strictly taxonomic (Dunbar & Shultz 2021a). Third, although some have claimed that phylogenetic methods should be used as a matter of course even where there is no signal, it is in fact bad statistical practice to include unnecessary random variables in an analysis: doing so necessarily reduces statistical power without adding explanatory value.

## Results

Fig. 2 plots mean species group size against residual neocortex volume (regressed on total brain volume). Although there is a significant overall correlation (OLS regression: r^2^=0.096, β=0.309, F_1,43_=4.55, p=0.039), the goodness of fit is poor and the data are clearly not bivariate normal. In such cases, RMA regression is recommended. An RMA regression yields:

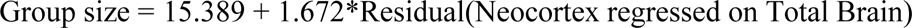

**Fig. 2.**
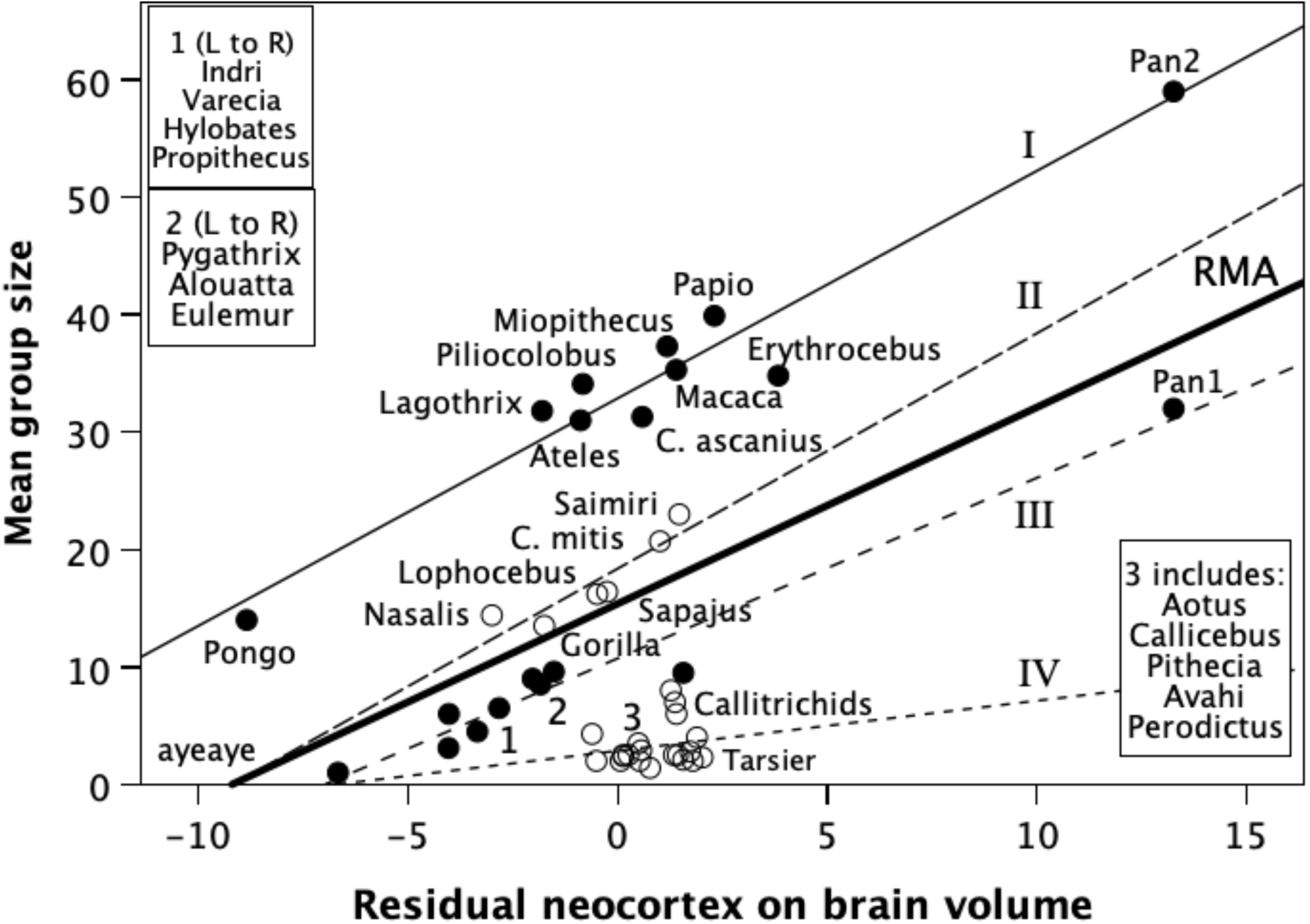
Mean genus social group size plotted against residual neocortex volume (regressed on total brain volume). Grades (identified by different symbols and Roman numerals) are determined by *k*-means cluster analysis of residuals from the RMA regression through the data (indicated by the thick regression line). The grade regression lines are OLS regressions. Two values are indicated for *Pan* corresponding to the bimodal peak (at N=32 and N=59) in the distribution of their community sizes (see Dunbar 2019). Values for two individual species are given for each of *Cercopithecus, Avahi* and *Cheirogaleus*: in each case, they lie on different grades.

with a marked improvement in goodness-of-fit (r^2^=0.749).

I therefore use the RMA regression (shown as the thick line) to calculate residual group size for each species (measured orthogonal to the regression line), and ran a *k*-means cluster analysis on these values to determine whether the data partition into natural groupings. The best fit is given by partitioning into *k*=4 clusters (Fig. 3), which a set of natural grades. An alternative would be a *k*=6 solution (Fig. 3), but, since all this does is to partition grades I and IV, parsimony would dictate *k*=4 as the better alternative.

**Fig. 3.**
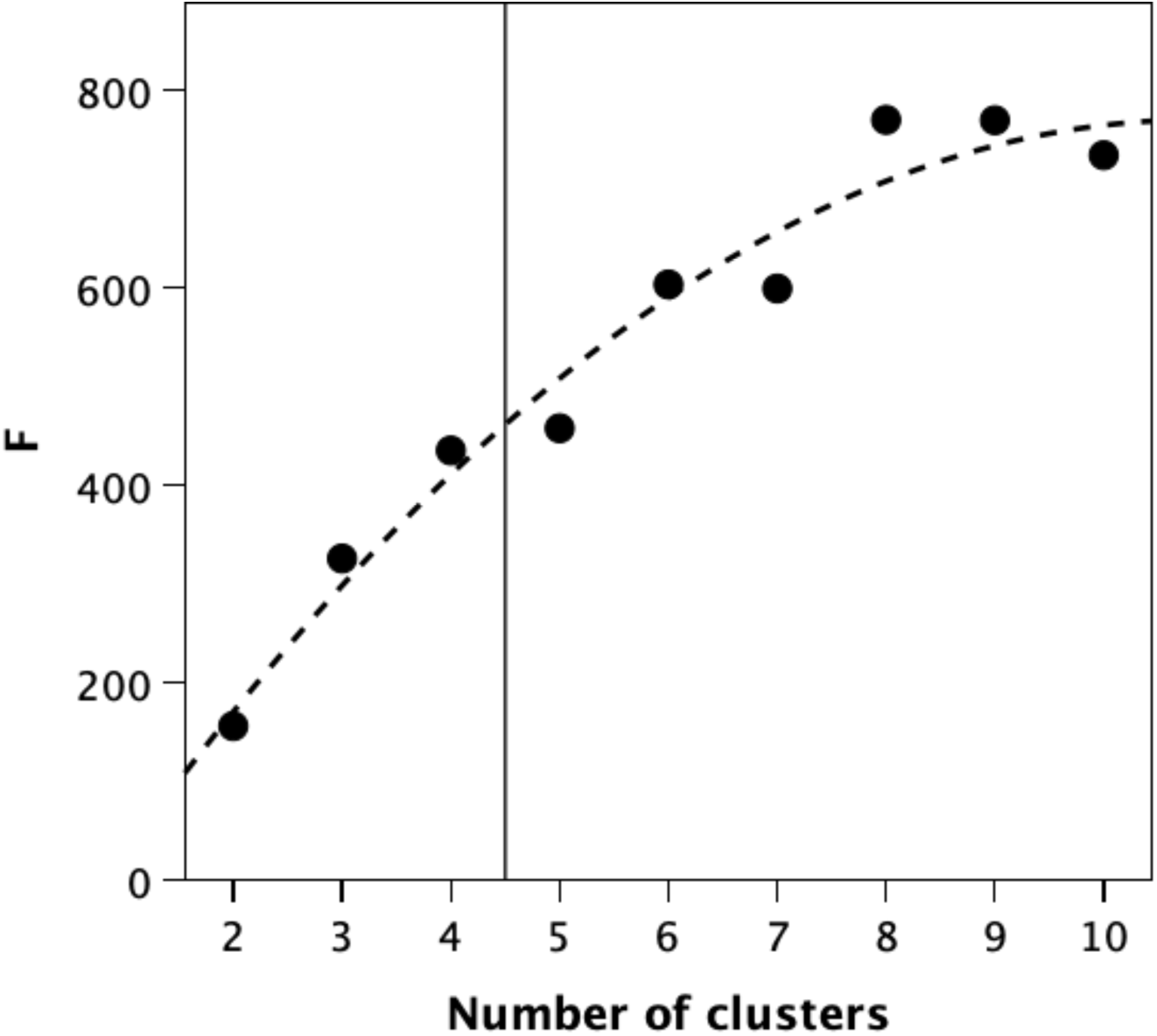
Goodness-of-fit (indexed as F-statistic) for *k-*means cluster analyses for 2≤*k*≤10. The optimal solution is the value of *k* where the F-static starts to asymptote, as defined by the cluster number 1/e^th^ down from the asymptote (indicated by the vertical line).

The uppermost cluster (grade I in Fig. 3) consists of a set of genera that either live in very large groups with some degree of fission-fusion flexibility (*Ateles*, *Lagothrix, Pan*), live in much larger groups than closely related genera of similar brain size (*Piliocolobus, Miopithecus, Erythrocebus, Cercopithecus ascanius*), or live in large stable, bonded groups (*Papio, Macaca*). Below them, grades II and III contain monkeys that live in relatively small, stable groups, plus *Gorilla* and *Hylobates* and some of the more social lemurines. These grades are distinguished mainly by group size, and a switch from harems to small multimale groups. Note that the two group sizes for *Pan* sit very close to the regression lines for grades I and III. Below these lies a cluster (grade IV) that consists of platyrrhines and strepsirrhines that live in monogamous pairs (of varying degrees of stability) and strepsirrhine genera that have a semi-solitary lifestyle. Note that the ayeaye is included in grade III despite its semi-solitary lifestyle, mainly because of its large negative neocortex residual.

Grade IV is notable for its division into two clear subclusters. The lefthand subcluster differs from the righthand subcluster both in relative neocortex size and in the greater stability of its groups. It includes a number of monogamous, pairbonded New World monkey genera (*Aotus, Pithecia, Callicebus*) that have very stable groups (longlasting pairbonds), as well as some Prosimians noted for having moderately stable groups (*Avahi, Lepilemur, Perodictus*). However, it also includes some prosimians that would usually be described as semi-solitary (*Otolemur, Nycticebus, Cheirogaleus*). The righthand subcluster consists entirely of genera that are semi-solitary (i.e. forage alone) even when they have a monogamous (or semi-monogamous) mating system and share sleeping nests. This subcluster includes tarsiers (an anthropoid), as well as the less social lemurines and galagines, and might legitimately include the callitrichids that sit directly above them to the extent that these have relatively unstable groups (Dunbar 1995a,b). The contrast between these two subclusters seems to be associated with a marked reduction in relative neocortex size (or, alternatively, an increase in subcortical brain volume).

The main distinction between the grades lies in group size. Mean group sizes for the four grades are 34.9, 17.4, 9.0 and 3.2, respectively. The differences are highly significant (F_3,42_=51.77, p<<0.0001), with most pairwise comparisons individually significant (Scheffé tests: I>II>III=IV, p<0.001). In effect, the grades correspond to a very clear progressive stepwise increase in group size. The transition from grade IV to grade III occurs at a group size of ∼5, and is associated with a decline in relative neocortex size. That between grades III and II occurs at groups of size 12-15 but is associated with an increase in relative neocortex size and demarcates a switch from small harems (grade III) to small multimale groups (grade II). That from grade II to grade I occurs at groups of 25-30 and is associated with a further increase in relative neocortex size.

I ran linear OLS regressions with three brain indices (neocortex volume, subcortical volume and total brain volume) as predictors of social group size for the full sample, first in separate univariate equations and then combined in alternative multivariate equations. I follow Kronmal’s (1993) advice and run the analysis with both the dominator variable and its reciprocal as a test of whether the outcome is determined by the numerator or the denominator. The results are given in Table 1. It is clear that although all three predictors individually yield significant regression equations, neocortex volume gives a better fit than either of the other two (models 1, 2A and 3A). More importantly, in the ratio version with neocortex as the numerator and either of the other variables as the denominator (models 2B and 3B) there is a significant fit for neocortex and a *nonsignificant* fit for both subcortex volume and total brain volume (with both the latter in any case having negative slopes).

**Table 1.** Regression analysis of the components of brain volume as predictors of group size.

| Model | Parameter estimates |  | df | t | p |
| --- | --- | --- | --- | --- | --- |
| | $\beta$ | $r^2$ | | | |
| **Univariate regressions** |  |  |  |  |  |
| Log10(Neocortex volume) | 0.703 | 0.494 | 43 | 6.48 | <0.0001 |
| Log10(Subcortical brain volume) | 0.629 | 0.395 | 43 | 5.30 | <0.0001 |
| Log10(Total brain volume) | 0.683 | 0.467 | 43 | 6.13 | <0.0001 |
| **Multiple regression model 1** [$r^2=0.630$, $F_{3,41}=35.71$, $p<<0.0001$] | | | | | |
| Log10(Neocortex volume) | -16.372 |  | 41 | -3.94 | 0.0003 |
| Log10(Subcortical volume) | -12.637 |  | 41 | -5.48 | <0.0001 |
| Log10(Total brain volume) | 29.558 |  | 41 | 4.63 | <0.0001 |
| **Multiple regression model 2A** [$r^2=0.500$, $F_{2,42}=20.97$, $p<<0.0001$] | | | | | |
| Log10(Neocortex volume) | 2.804 |  | 42 | 5.16 | <0.0001 |
| Log10(Subcortical volume) | -2.134 |  | 42 | -3.93 | <0.001 |
| **Multiple regression model 2B** [$r^2=0.500$, $F_{2,42}=20.97$, $p<<0.0001$] | | | | | |
| Log10(Neocortex volume) | 0.715 |  | 42 | 6.47 | <0.0001 |
| 1/(Log10Subcortical volume) | -0.077 |  | 42 | -0.70 | 0.491 |
| **Multiple regression model 3A** [$r^2=0.579$, $F_{2,42}=28.93$, $p<<0.0001$] | | | | | |
| Log10(Neocortex volume) | 5.380 |  | 42 | 3.36 | 0.002 |
| Log10(Brain volume) | -4.686 |  | 42 | -2.92 | 0.006 |
| **Multiple regression model 3B** [$r^2=0.512$, $F_{2,42}=20.97$, $p<<0.0001$] | | | | | |
| Log10(Neocortex volume) | 0.863 |  | 42 | 5.13 | <0.0001 |
| 1/Log10(Brain volume) | 0.209 |  | 42 | 1.24 | 0.221 |

**Table 2.** Multiple regression of predictors of cerebellum volume.

| Model | Parameter estimate |  | df | t | p |
| --- | --- | --- | --- | --- | --- |
| | $\beta$ | $r^2$ | | | |
| **Cerebellum volume (cc)** [$r^2=0.976$, $F_{3,18}=245.67$, $p<<0.0001$] | | | | | |
| Body mass (kg) | 0.965 |  | 18 | 24.2 | <0.001 |
| Diet (% fruit) | 0.092 |  | 18 | 2.19 | 0.042 |
| Group size | 0.095 |  | 18 | 2.20 | 0.042 |
| **Cerebellum/neocortex (%)** [$r^2=0.518$, $F_{3,15}=5.38$, $p=0.010$] | | | | | |
| Body mass (kg) | 0.275 |  | 15 | 1.42 | 0.177 |
| Diet (% fruit) | -0.179 |  | 15 | -0.91 | 0.380 |
| Group size | -0.589 |  | 15 | -3.22 | 0.006 |

This is reflected in the relationship between neocortex volume and the volume of the subcortical brain across the grades (Fig. 4). Fig. 4a suggests that, as genera have evolved across the grades, subcortical brain size has become relatively smaller while neocortex has become progressively larger. In addition, although the regressions for the four grades are individually significant (grade I: r^2^=0.955, β=0.977, F_1,8_ = 170.6, p<<0.0001; grade II: r^2^=0.998, β=0.999, F_1,4_ = 10036.7, p<<0.0001; grade III: r^2^=0.931, β=0.965, F_1,8_ = 108.2, p<0.0001; grade IVA: r^2^=0.925, β=0.962, F_1,8_ = 99.0, p<<0.0001; grade IVB: r^2^=0.831, β=0.911, F_1,6_ = 29.4, p=0.002), there is a progressive decrease in slope across the grades (Spearman correlation: r_S_=0.900, df=5, p=0.037). In other words, at each phase change in group size grade, increasing investment is being made in neocortical tissue, without necessarily reducing the investment in subcortical tissue. Fig. 4b suggests that this is, in part at least, associated with an increase in body mass: the overall regression changes pitch at around 0.6 kg, 3 kg and 6 kg body mass. Initially, subcortical brain volume increases roughly linearly with body mass up to about 2 kg body mass, and then rapidly levels off with increases in brain mass being invested disproportionately in neocortex volume. This pattern applies across the taxonomic groups.

**Fig. 4.**
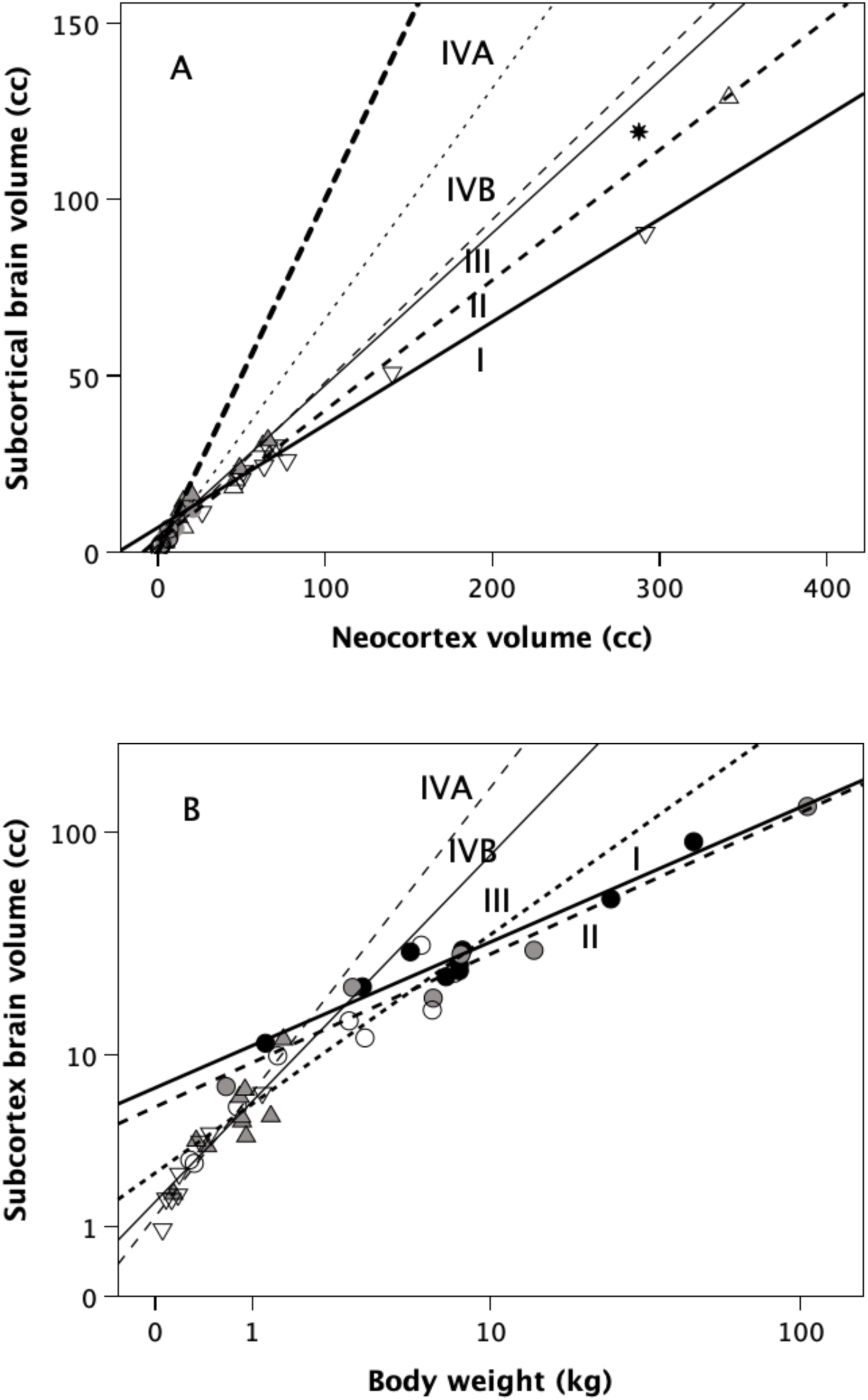
(a) Body mass and (b) neocortex volume for pairliving and semi-solitary genera in grades III and IV. A and B distinguish the left and right subclusters in grade IV (Fig. 2). The bars in (a) indicate the mean for each group.

Fig. 5 suggests that the switches in Fig. 4 are associated mainly with locomotor style. The cerebellum:neocortex ratio (expressed as a percentage) is high in prosimians (including tarsiers), all of whom are either clingers-and-leapers or slow arboreal quadrupeds, but low in arboreal anthropoids and very low in terrestrial anthropoids (with the exception of the gorilla). The gorilla’s high cerebellum ratio (and, to a lesser extent, that of chimpanzees) may be associated with the fact that both are partially arboreal, suggesting that arboreality may be especially demanding for a large-bodied animal. In this, they contrast with *Homo* (star symbol) who sit with the most terrestrial anthropoids (*Erythrocebus, Papio, Macaca*). Note that the brachiators (*Hylobates, Ateles*) and semi-brachiators (*Nasalis, Lagothrix*) have a high cerebellum ratio, suggesting that the cerebellum plays an important role in their unusual locomotory style.

**Fig. 5.**
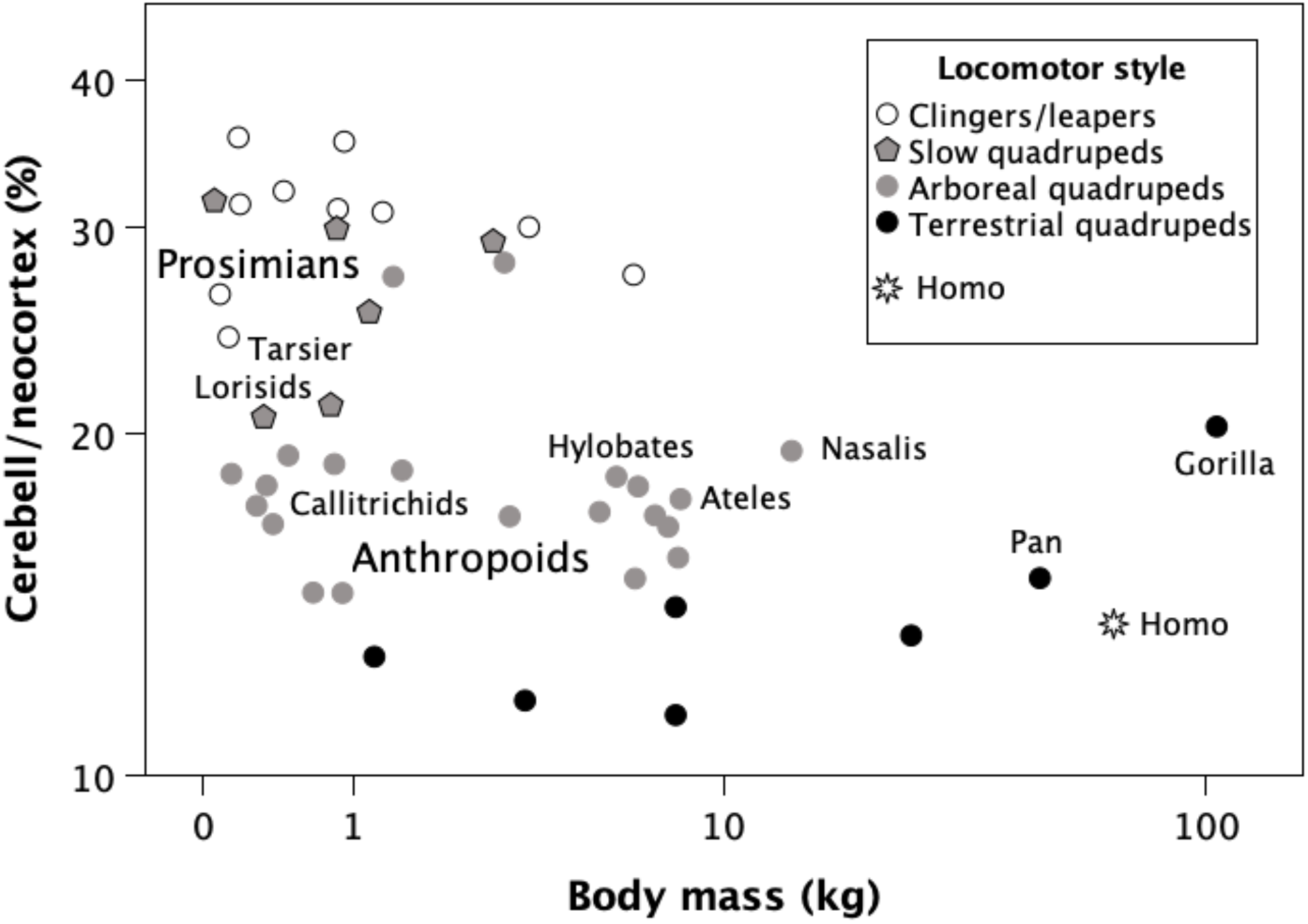
Subcortical brain volume as a function of (a) neocortex volume and (b) body mass, for the four grades shown in Fig. 3. Grades are indicated by roman numerals. Grade IV is partitioned into (A) the righthand subcluster and (B) the lefthand subcluster. The heavy-dashed line on the left is the line of equality (subcortex volume = neocortex volume).

**Fig. 5.** Cerebellum:neocortex ratio (indexed as a percentage) in primate genera as a function of body mass and locomotor style. Star: *Homo*.

## Discussion

The relative investment in brain regions associated with functional cognition (as opposed to somatic tissue management) varies across primates in ways that correlate with group size and demographic structure. This does not involve a single parametric relationship, but rather a series of grades that constitute a stepwise pattern in which there is a phase shift in group size at each transition point. These phase shifts seem to coincide with the points at which novel socio-cognitive strategies designed to engineer group cohesion are introduced (Dunbar 2025). The progressive increase in relative neocortex size in grades III through I suggests that managing large groups may be cognitively increasingly demanding as groups get larger (see also Dunbar 2025).

Although it has been suggested that the cerebellum might play an important role in sociality, this seems not to be the case: Fig. 5 suggests that the switches to higher demographic grades are associated with a *reduction* in absolute cerebellum size. A terrestrial lifestyle typically incurs a significantly higher level of predation risk (Hill & Lee 1994; Dunbar & Shultz 2021b), necessitating an increase in group size as a defence. However, a terrestrial lifestyle involves fewer complex locomotory decisions, and this may allow investment in the cerebellum to be reduced relative to that in the neocortex.

The position of the pairliving genera is noteworthy for two reasons. One is that they form a distinct grade of their own (grade IV, with some anthropoids on grade III); the other is that these pairliving genera are split across three separate subclusters. Aside from the callitrichids, all of the genera in grade IVB are nocturnal, whereas most (but not all) of the genera in grade IVA and the base of grade III are diurnal, with the latter having relatively smaller neocortices than the former. This is unlikely to be due to differences in visual system, since the diurnal species have visual system volumes that are, as a percentage of both total brain and neocortex volume, significantly larger due to the high computational costs of colour vision (Barton 1996).

What does differentiate the six pairliving genera on grade III is that, like the gorilla, they are the largest-bodied members of their respective taxonomic subfamilies. In addition, the gibbon is a brachiator who, like the semi-brachiators *Ateles* and *Lagothrix* and the quadrupedal *Nasalis*, are large-bodied and have relatively large cerebella, suggesting that the coordination demands of manoeuvring large bodies in three-dimensional space in trees may be more cognitively demanding. So the small relative sizes of their neocortices may simply be due to the relatively large size of their cerebella (Fig. 5). Taken together, these results suggest that the cerebellum (one of whose crucial functions is motor coordination) is more important functionally for arboreal taxa generally, and especially so those genera that engage in distance jumping and suspensory brachiation. It is much less necessary for a terrestrial lifestyle, allowing more space to be devoted to the neocortex, and hence larger groups.

